# RNAseqCovarImpute: a multiple imputation procedure that outperforms complete case and single imputation differential expression analysis

**DOI:** 10.1101/2023.05.11.540260

**Authors:** Brennan H. Baker, Sheela Sathyanarayana, Adam A. Szpiro, James MacDonald, Alison G. Paquette

## Abstract

Missing covariate data is a common problem that has not been addressed in observational studies of gene expression. Here we present a multiple imputation (MI) method that accommodates high dimensional transcriptomic data by binning genes, creating separate MI datasets and differential expression models within each bin, and pooling results with Rubin’s rules. Simulation studies using real and synthetic data show that this method outperforms complete case and single imputation analyses at uncovering true positive differentially expressed genes, limiting false discovery rates, and minimizing bias. This method is easily implemented via an R package, “RNAseqCovarImpute” that integrates with the limma-voom pipeline.

## Background

Missing data is a common problem in observational studies, as modeling techniques such as linear regression cannot be fit to data with missing points. Missing data is frequently handled using complete case analyses in which any individuals with missing data are dropped from the study. Dropping participants can reduce statistical power and, in some cases, result in biased model estimates. A common technique to address these problems is to replace or ‘impute’ missing data points with substituted values. Typically, for a given covariate, missing data points are imputed using a prediction model including other relevant covariates as independent variables. In single imputation, a missing value is replaced with the most likely value based on the predictive model. However, by ignoring the uncertainty inherent with predicting missing data, single imputation methods can result in biased coefficients and over-confident standard errors (1). Multiple imputation addresses this problem by generating several predictions, thereby allowing for uncertainty about the missing data. In a typical multiple imputation procedure: 1) M imputed data sets are created, 2) each data set is analyzed separately (e.g., using linear regression), and 3) estimates and standard errors across the M analyses are pooled using Rubin’s rules (2, 3).

To date, there has been no concerted effort to determine the most advantageous method for handling missing covariate data in transcriptomic studies. A large proportion of RNA-sequencing (RNA-seq) studies are conducted in in-vitro or in vivo models and do not suffer from missing covariate data. Complete datasets are common in experimental studies with controlled conditions and a limited number of covariates. In an experimental setting, studies may employ two-group analyses with no additional variables or utilize covariates for which collecting data is trivial (e.g., sequencing batch and sex). However, the cost of sequencing has decreased over time (4), and transcriptomic data are already becoming more common in large human observational studies where missing data is a prevailing concern (5, 6). Therefore, guidelines for handling missing data in this context are critically needed to facilitate the integration of transcriptomic and epidemiologic approaches.

While single imputation methods must omit the outcome from the imputation predictive model to avoid bias, the opposite is true of multiple imputation (7). However, including the outcome in the multiple imputation predictive model can be problematic in ‘omics’ studies with high dimensional data. Fitting an imputation model where the number of independent variables is far greater than the number of individuals in the study is generally not feasible. For instance, in RNA-seq studies with tens of thousands of genes, an equal or greater number of participants may be needed to apply a standard multiple imputation procedure.

One solution is to make one set of M imputed datasets per gene, where expression data for a single gene is included in the predictive model. Then, each set of imputed data can be used to estimate differential expression of the gene that was used in that set’s predictive modeling. However, the generation of tens of thousands of sets of imputed data is computationally intensive. In epigenetic studies of DNA methylation at CpG cites, this approach has been modified to be less computationally intensive by using groups of CpG sites together to impute missing data (8, 9). Here, we developed a similar method that bins genes into smaller groups for imputing missing values of covariates in RNA-seq studies. We created an R package (RNAseqCovarImpute) that is fully compatible with the popular limma-voom (10-12) differential expression analysis pipeline. We conducted a simulation study to compare the performance of multiple imputation as implemented in RNAseqCovarImpute with random forest single imputation and complete case analyses. Finally, we applied RNAseqCovarImpute to an analysis of placental transcriptomic changes associated with maternal age.

## Results

### Multiple imputation and differential expression analysis in RNAseqCovarImpute package

The RNAseqCovarImpute package implements multiple imputation of missing covariates and differential gene expression analysis by 1) randomly binning genes into smaller groups, 2) creating M imputed datasets separately within each bin, where the imputation predictor matrix includes all covariates and the log counts per million (CPM) for the genes within each bin, 3) estimating gene expression changes using voom (10) followed by lmFit (11, 12) functions, separately on each M imputed dataset within each gene bin, 4) un-binning the gene sets and stacking the M sets of model results before applying the squeezeVar (11, 12) function to apply a variance shrinking Bayesian procedure to each M set of model results, 5) pooling the results with Rubins’ rules to produce combined coefficients, standard errors, and P-values, and 6) adjusting P-values for multiplicity to account for false discovery rate (FDR) (13). An overview of how the method would be applied to a study measuring expression for 10,000 genes in 500 individuals/observations is shown in Figure 1.

**Figure 1:**
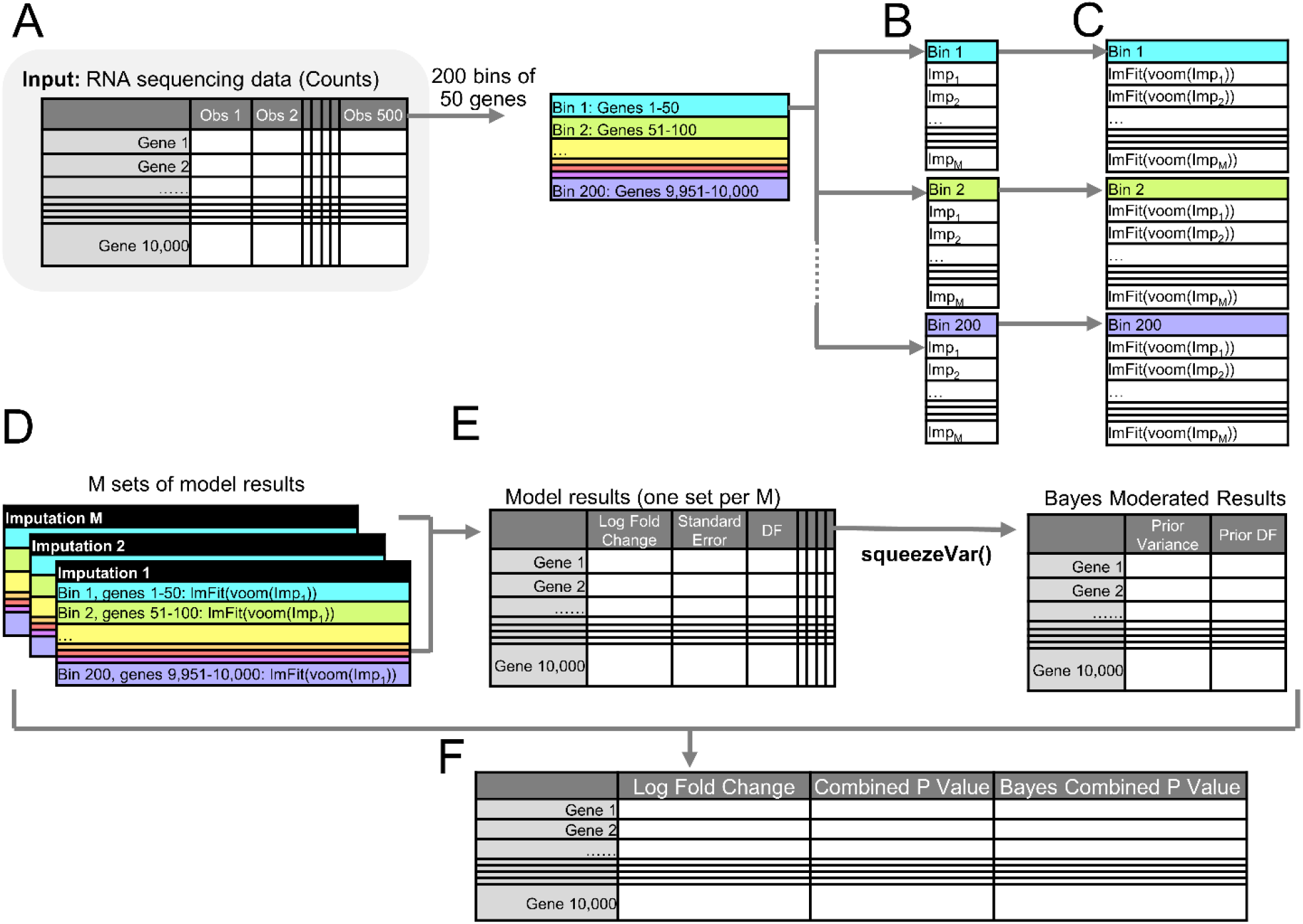
Overview of Multiple Imputation Method Legend: A) Starting with a matrix of counts for 10,000 genes across 500 observations, create 200 bins of 50 genes (default = 1 gene per 10 observations). B) Separately within each bin, create M imputed datasets (Imp_1_ – Imp_m_) using mice function. Along with covariates, the imputation predictor matrix includes the log-CPM values for the genes within each bin. C) Estimate association of predictor of interest with gene expression using voom followed by lmFit functions. These functions are run separately within each gene bin on each M imputed dataset created using those genes in the predictor matrix. Rather than calculating precision weights separately within each bin, a modified voom function utilizes the mean-variance curve fit using all genes across all M imputations. D) Un-bin genes and stack into M sets of model results. E) Use squeezeVar function separately on each M set of model results to squeeze variances using an empirical Bayes procedure. F) Separately for the original and Bayes modified results, combine across M sets of model results using Rubin’s rules. Adjust p-values for multiplicity using false discovery rate (FDR).

### Simulation studies

We performed three separate simulation studies to compare multiple imputation implemented via RNAseqCovarImpute with single imputation (SI) and complete case (CC) analyses. Simulations were conducted using 1) real covariate and real RNA-seq data from 1,044 individuals, 2) real covariate data and synthetic RNA-seq data with known signal, and 3) synthetic covariate data with known confounding structure and synthetic RNA-seq data with known signal (see methods for details). In all three simulation studies, we first conducted differential expression analysis using the limma-voom pipeline on the entire set of observations with their complete covariate data (hereinafter “full data”). These models estimated the effect of a predictor of interest on gene expression while controlling for several covariates. Genes significantly associated with the predictor of interest at FDR <0.05 in these full data models were defined as true differentially expressed genes (DEGs). Missingness was then simulated to emulate a common situation in scientific research where an investigator has complete data for a predictor of interest, but may have missing data for other important covariates. Therefore, missingness was only induced in covariates and not the predictor of interest. We explored scenarios with various levels of missing data ranging from 5-55% of participants having at least one missing data point, and under two missingness mechanisms: missing completely at random (MCAR) and missing at random (MAR). We simulated ten datasets for each missingness mechanism at each level of missingness before applying RNAseqCovarImpute, SI, and CC methods and comparing the results with the full data model. Missingness was simulated using the ampute function from the mice package (14). CC analyses dropped any individual with at least one missing data point, while SI imputed missing data using the missForest package (15). The limma-voom pipeline was applied for CC and SI as described for the full data model.

Our objective was to evaluate the ability of RNAseqCovarImpute, SI, and CC methods to identify true DEGs from the full data model as significant while limiting false positives. True DEGs from the full data model that were also identified as significant by a given method were defined as true positives. We report the percent of true DEGs identified as significant for each method out of the total number of true DEGs from the full data model. Genes erroneously identified as significant by a given method that were not true DEGs from the full data analysis were defined as false positives. We report the percent of false positives out of the total number of significant results for each method as the FDR. Bias was calculated as the difference between coefficients from the full data model and coefficients produced by each method, expressed as a percent of the full data model coefficient. We report the average bias per gene per missingness scenario across all 10 simulations.

### Simulation #1: real covariates and real counts

The full data model uncovered 2,453 true DEGs associated with the predictor of interest while controlling for covariates. When simulated covariate missingness was MCAR, both imputation methods identified more true DEGs than CC (Figure 2A). RNAseqCovarImpute identified a larger percent of true DEGs than SI across all levels of missingness (range = 95.6-98.5% and 56.6-90.7%, respectively). RNAseqCovarImpute, SI, and CC methods performed similarly with respect to FDR (range = 2.9-13.8%, 1.9-14.8%, and 5.3-12.8%, respectively). Bias was more tightly centered around zero for RNAseqCovarImpute with respect to all genes (Figure 2B) and when only the 2,453 true DEGs were included (Figure 2C).

**Figure 2:**
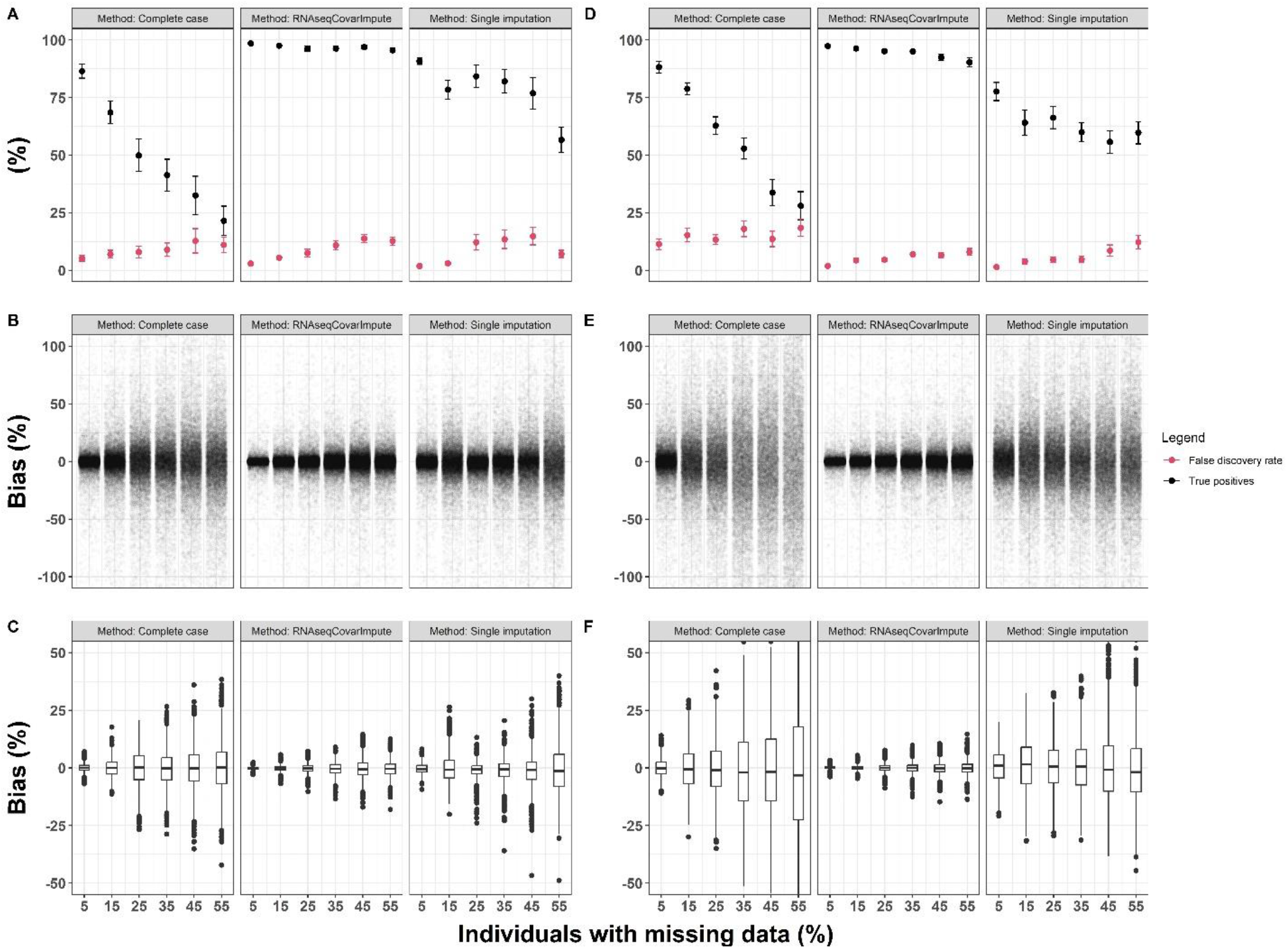
Simulation #1: Real Covariates and Real CountsLegend: Performance of complete-case, RNAseqCovarImpute, and single imputation analyses on ten datasets with simulated missingness per missingness mechanism and level of missingness versus the full data differential expression model on the real covariate and RNA-seq data (N = 1,044). Error, bias for all genes, and bias for only true DEGs shown, respectively, for MCAR (A, B, C) and MAR (D, E, F) missingness mechanisms. Mean ± standard error shown for false discovery rate (red) and percent of true positives identified (black). Average bias shown for each gene as a percent of the full data model differential expression coefficient. Bias for all genes shown with one point per gene (B, E), while bias for true DEGs shown with box (median and interquartile range) and whisker (1.5* interquartile range) plots (C, F).

When data were MAR, both imputation methods identified more true DEGs and fewer false positives compared to CC (Figure 2D). Again, RNAseqCovarImpute identified a larger percent of true DEGs than SI across all levels of missingness (range = 90.3-97.4% and 55.7-77.6%, respectively). While RNAseqCovarImpute and SI performed similarly with respect to FDR (range = 2.0-8.0% and 1.4-12.2%, respectively), FDR for CC was substantially higher (11.4-18.5%; Figure 2D). Again, bias was more tightly centered around zero for RNAseqCovarImpute with respect to all genes (Figure 2E) and when only true DEGs were included (Figure 2F).

### Simulation #2: real covariates and synthetic counts

RNA-seq data were modified to add known signal using the seqgendiff package (16). As an initial diagnostic, we confirmed that the coefficients for each gene estimated from the limma-voom pipeline on this synthetic count table with no missingness closely matched our user defined coefficients (Supplemental Figure 1). In the set of simulations with a null gene association rate of 82.5% and an MCAR missingness mechanism, RNAseqCovarImpute identified the highest percent of true DEGs, followed sequentially by SI and CC (range = 96.8-99.1%, 78.9-95.9%, and 58.2-92.9%, respectively; Figure 3A). With respect to FDR, RNAseqCovarImpute and SI performed similarly (range 1.4-6.1% and 1.4-7.6%, respectively), while higher FDRs were observed for CC (range = 3.0-8.0%; Figure 3A). Bias was more tightly centered around zero for RNAseqCovarImpute compared to SI and CC (Figure 3B,C) Patterns were similar when data were MAR with respect to error (RNAseqCovarImpute, SI, and CC percent true DEGs ranges = 94.0-98.2%, 77.6-89.6%, and 61.1-93.2%, and FDR ranges = 0.9-3.9%, 1.2-7.7%, and 6.0-10.9%, respectively; Figure 3D) and bias (Figure 3E,F).

**Figure 3:**
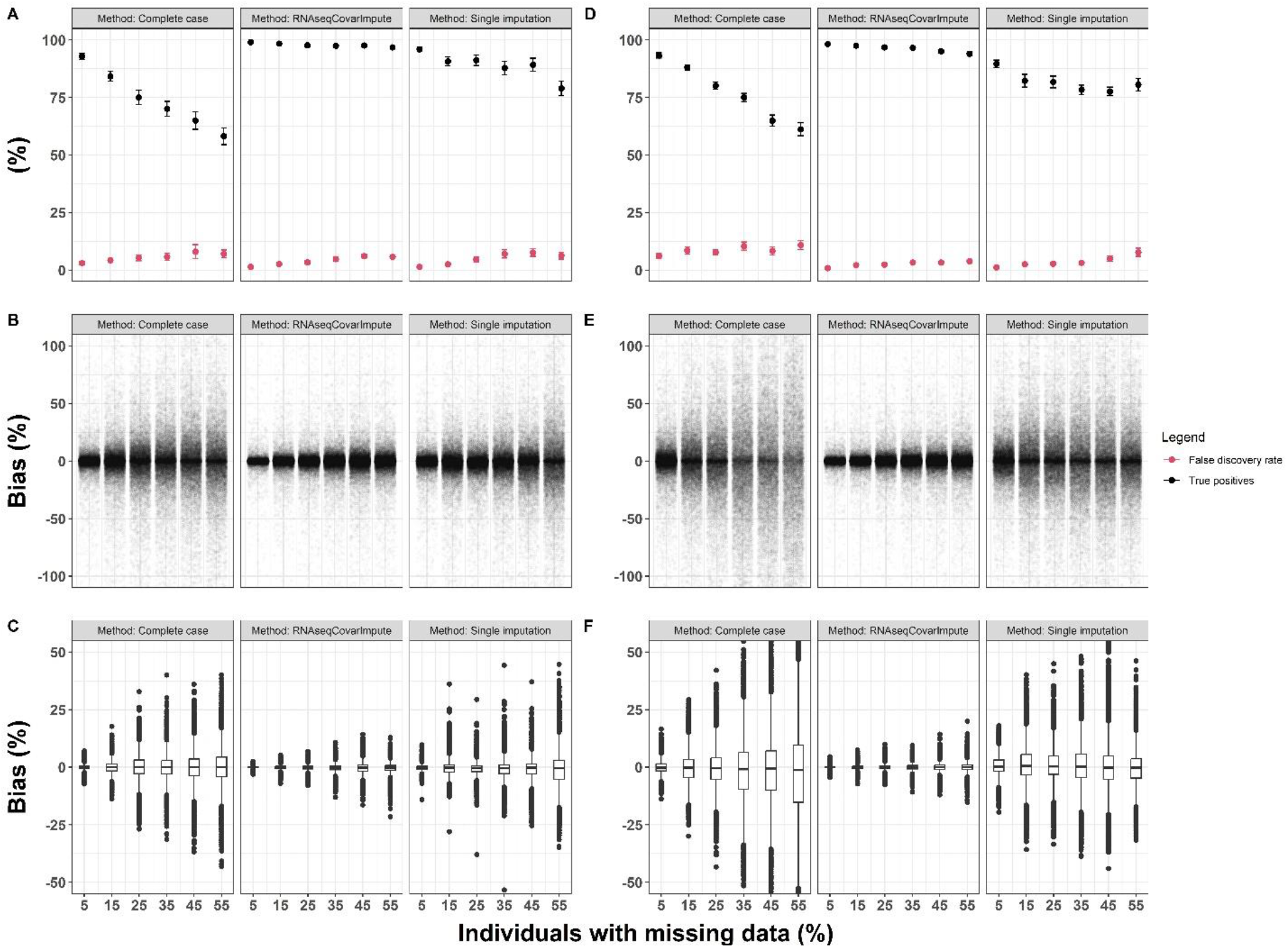
Simulation #2: Real Covariates and Synthetic Counts with 82.5% Null Gene RateLegend: Performance of complete-case, RNAseqCovarImpute, and single imputation analyses on ten datasets with simulated missingness per missingness mechanism and level of missingness versus the full data differential expression model on the real covariate and synthetic RNA-seq data with 82.5% null gene association rate (N = 1,044). Error, bias for all genes, and bias for only true DEGs shown, respectively, for MCAR (A, B, C) and MAR (D, E, F) missingness mechanisms. Mean ± standard error shown for false discovery rate (red) and percent of true positives identified (black). Average bias shown for each gene as a percent of the full data model differential expression coefficient. Bias for all genes shown with one point per gene (B, E), while bias for true DEGs shown with box (median and interquartile range) and whisker (1.5* interquartile range) plots (C, F).

All three methods performed slightly better in our sensitivity analysis with a null gene association rate of 50% rather than 82.5%, with similar patterns when we compared the methods. In general, RNAseqCovarImpute performed better than SI, and SI performed better than CC with respect to identifying true DEGs, limiting FDR, and limiting bias of true DEG coefficients under both MCAR (Figure 4A-C) and MAR (Figure 4D-F) missingness mechanisms.

**Figure 4:**
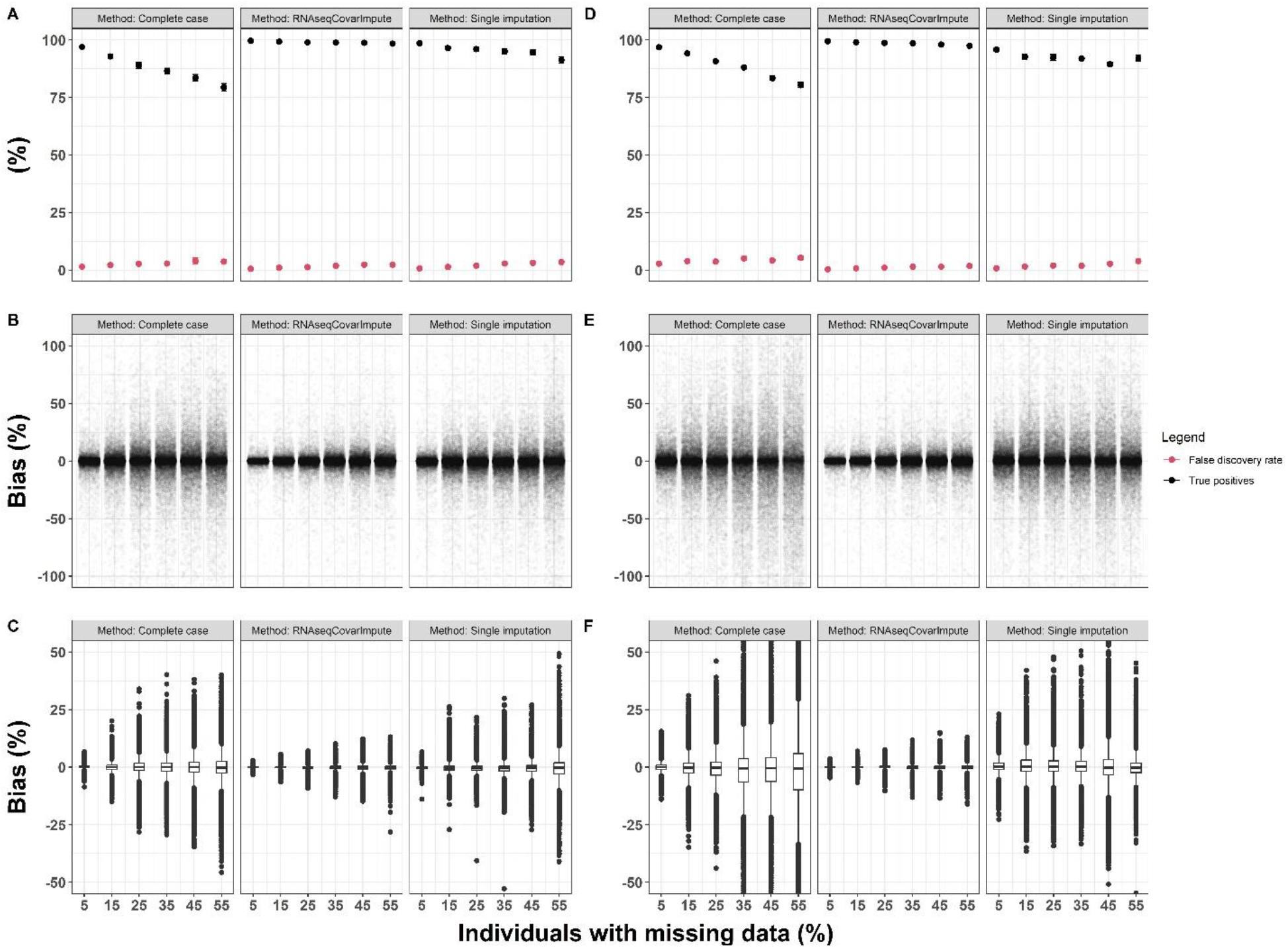
Simulation #2: Real Covariates and Synthetic Counts with 50% Null Gene RateLegend: Performance of complete-case, RNAseqCovarImpute, and single imputation analyses on ten datasets with simulated missingness per missingness mechanism and level of missingness versus the full data differential expression model on the real covariate and synthetic RNA-seq data with 50% null gene association rate (N = 1,044). Error, bias for all genes, and bias for only true DEGs shown, respectively, for MCAR (A, B, C) and MAR (D, E, F) missingness mechanisms. Mean ± standard error shown for false discovery rate (red) and percent of true positives identified (black). Average bias shown for each gene as a percent of the full data model differential expression coefficient. Bias for all genes shown with one point per gene (B, E), while bias for true DEGs shown with box (median and interquartile range) and whisker (1.5* interquartile range) plots (C, F).

### Simulation #3: synthetic covariates and synthetic counts

Synthetic covariates were generated with a goal of V1 (the predictor of interest) being correlated with V2-V5 (confounders) at Spearman’s rank correlation coefficient of 0.1, which was reasonably well reflected in the realized correlations (Supplemental Figure 2). In general, RNAseqCovarImpute was the best performing method at identifying true DEGs and limiting FDR across all sample sizes (Figure 5). Moreover, across all missingness mechanisms, sample sizes, and levels of missingness, bias was more tightly centered around 0 for RNAseqCovarImpute compared with CC and SI methods with respect to all genes (Figure 6) and when only true DEGs were included (Figure 7). In sensitivity analyses exploring different correlations of V1 with V2-V5, RNAseqCovarImpute remained the best performer in general (Supplemental Figure 3). However, when covariate correlations with V1 were set to 40%, SI identified slightly more true positives than RNAseqCovarImpute, but at the cost of substantially higher FDR (FDR>60% in one instance; Supplemental Figure 3B).

**Figure 5:**
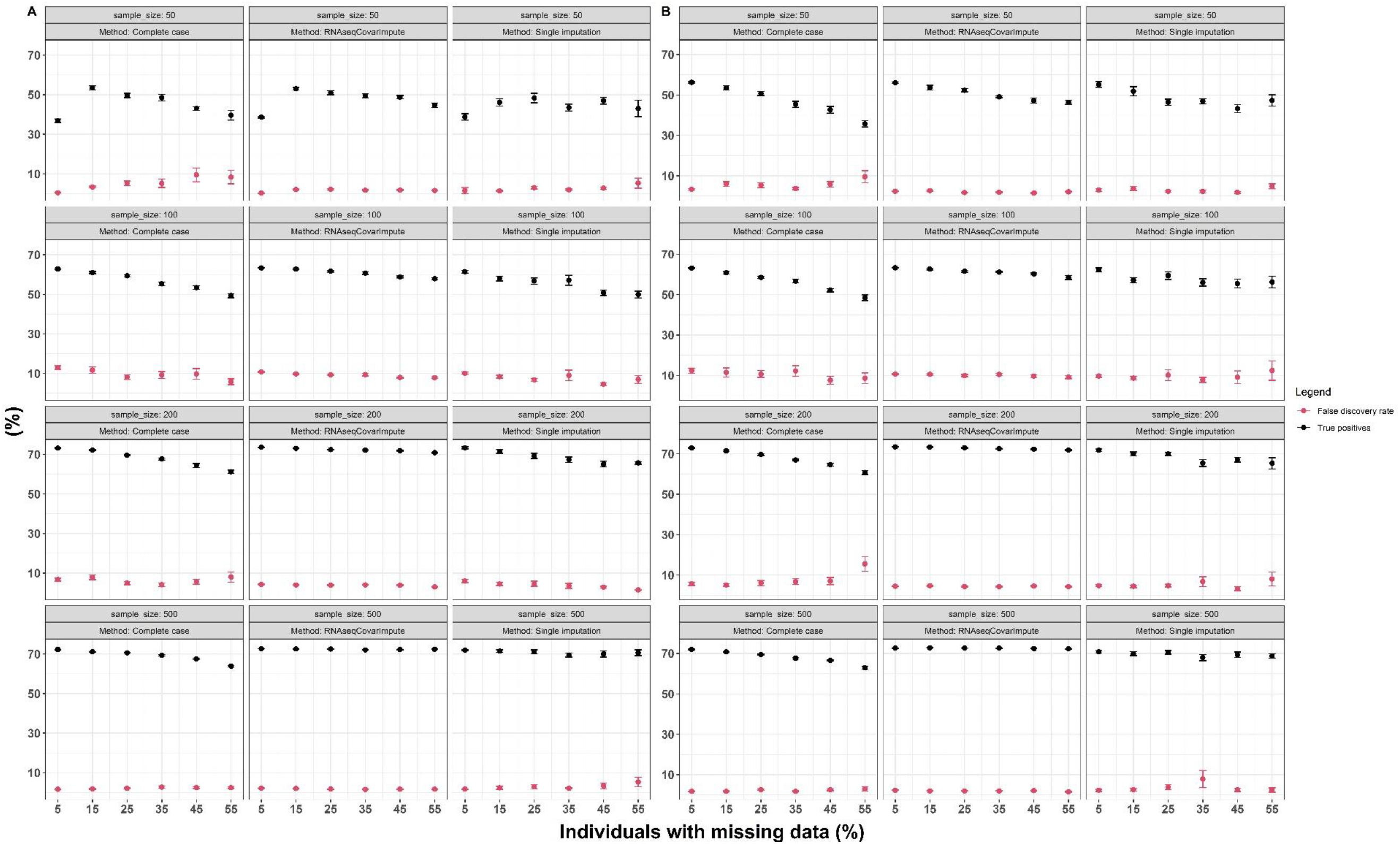
Simulation #3: Synthetic Covariates and Synthetic Counts ErrorLegend: Performance of complete-case, RNAseqCovarImpute, and single imputation analyses on ten datasets with simulated missingness per missingness mechanism, level of missingness, and sample size versus the full data differential expression model on the fully synthetic covariate and RNA-seq data. Mean ± standard error shown for false discovery rate (red) and percent of true positives identified (black) for MCAR (A) and MAR (B) missingness mechanisms.

**Figure 6:**
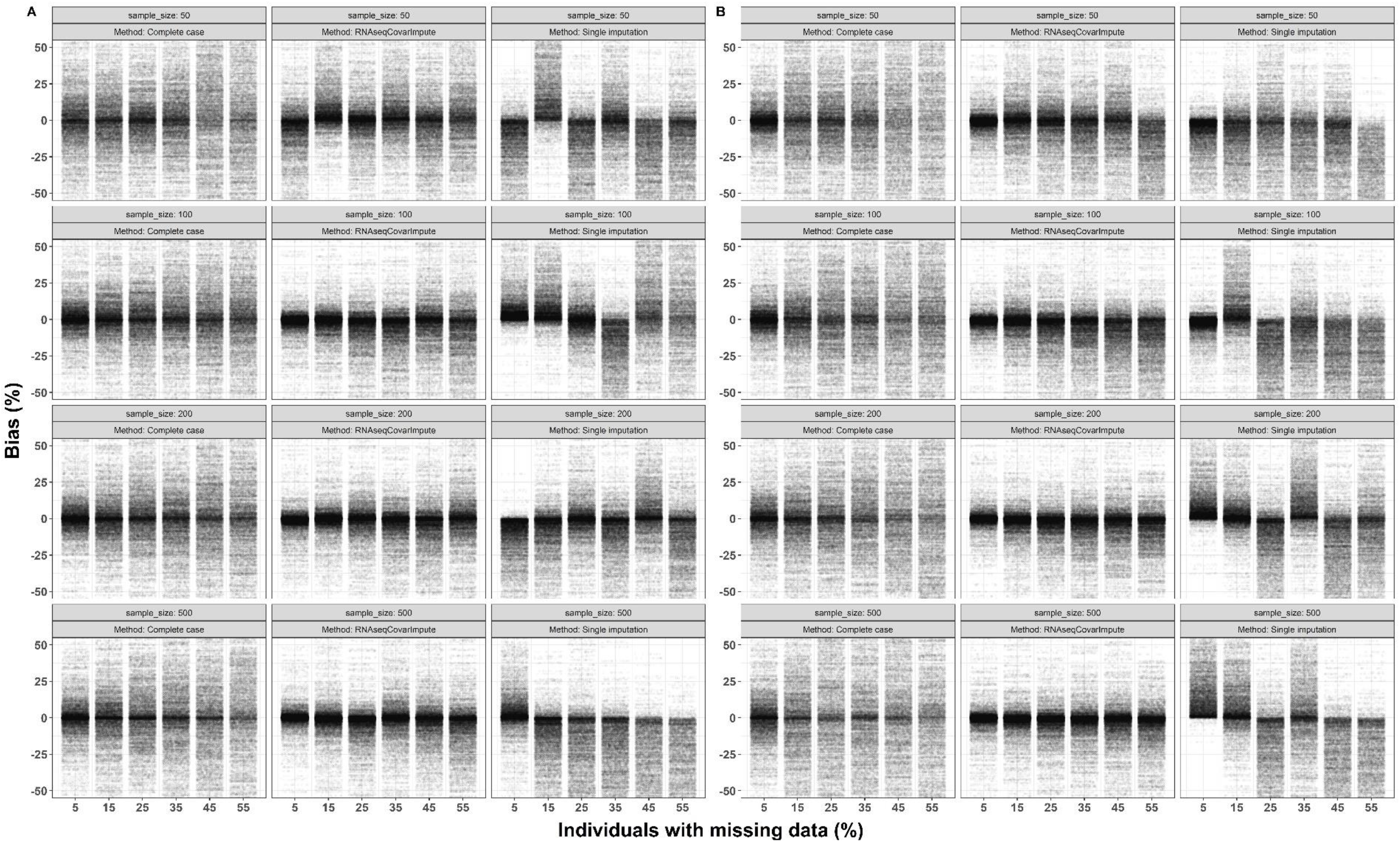
Simulation #3: Synthetic Covariates and Synthetic Counts BiasLegend: Performance of complete-case, RNAseqCovarImpute, and single imputation analyses on ten datasets with simulated missingness per missingness mechanism and level of missingness versus the full data differential expression model on the fully synthetic covariate and RNA-seq data. Average bias shown for each gene as a percent of the full data model differential expression coefficient. Bias for all genes shown with one point per gene for MCAR (A) and MAR (B) missingness mechanisms.

**Figure 7:**
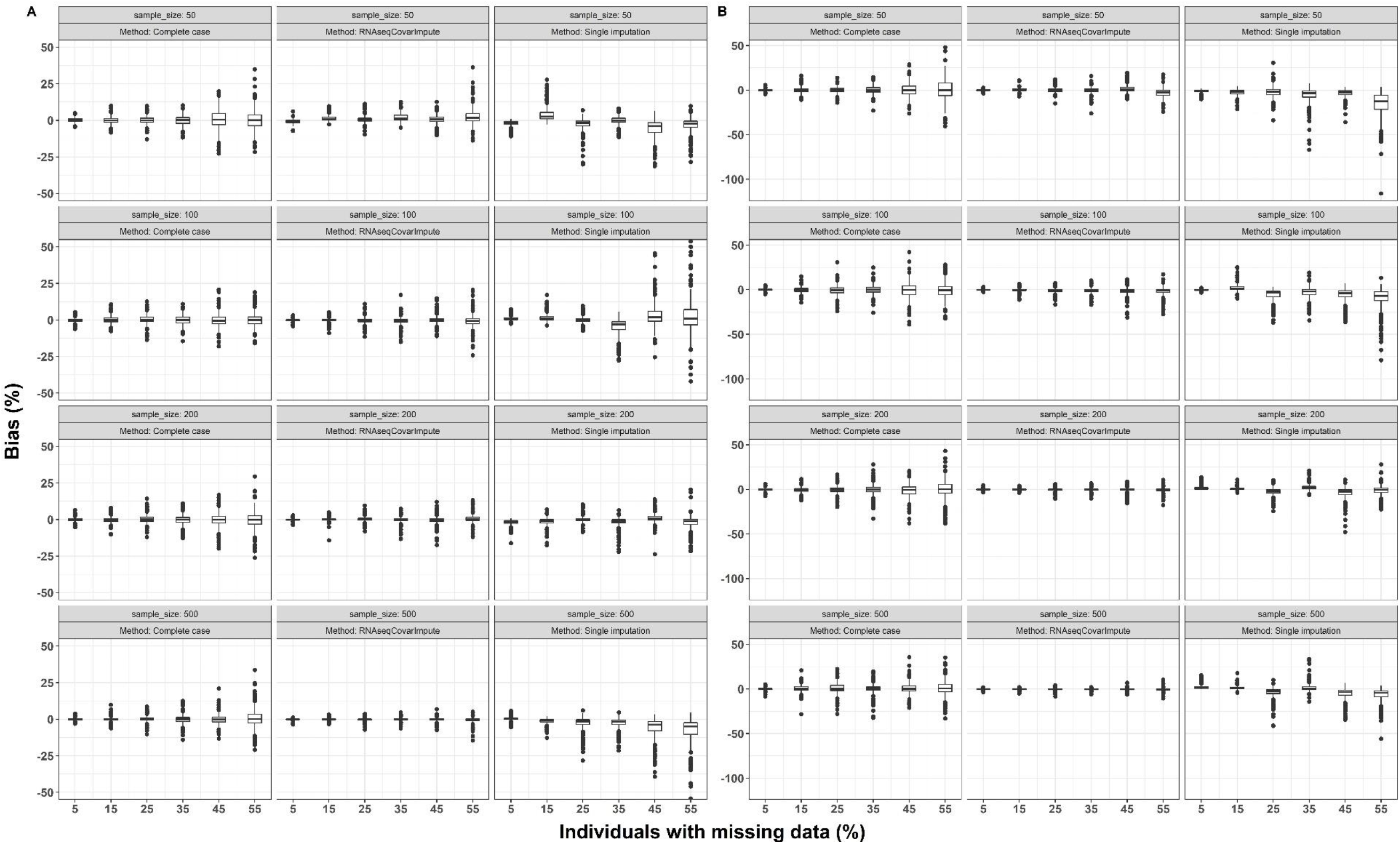
Simulation #3: Synthetic Covariates and Synthetic Counts BiasLegend: Performance of complete-case, RNAseqCovarImpute, and single imputation analyses on ten datasets with simulated missingness per missingness mechanism and level of missingness versus the full data differential expression model on the fully synthetic covariate and RNA-seq data. Bias for true DEGs shown with box (median and interquartile range) and whisker (1.5* interquartile range) plots for MCAR (A) and MAR (B) missingness mechanisms.

### Application of RNAseqCovarImpute in analysis of maternal age and placental transcriptome

To demonstrate how RNAseqCovarImpute can implement multiple imputation and the limma-voom pipeline in a real-world example, we applied it to the largest placental transcriptomic dataset to-date, which was generated by the ECHO prenatal and early childhood pathways to health (ECHO-PATHWAYS) consortium (5). This analysis examined the association of maternal age with the placental transcriptome while controlling for 10 covariates including potential confounders, mediators, and precision variables. The causal relationships among these variables are illustrated in Figure 8.

**Figure 8:**
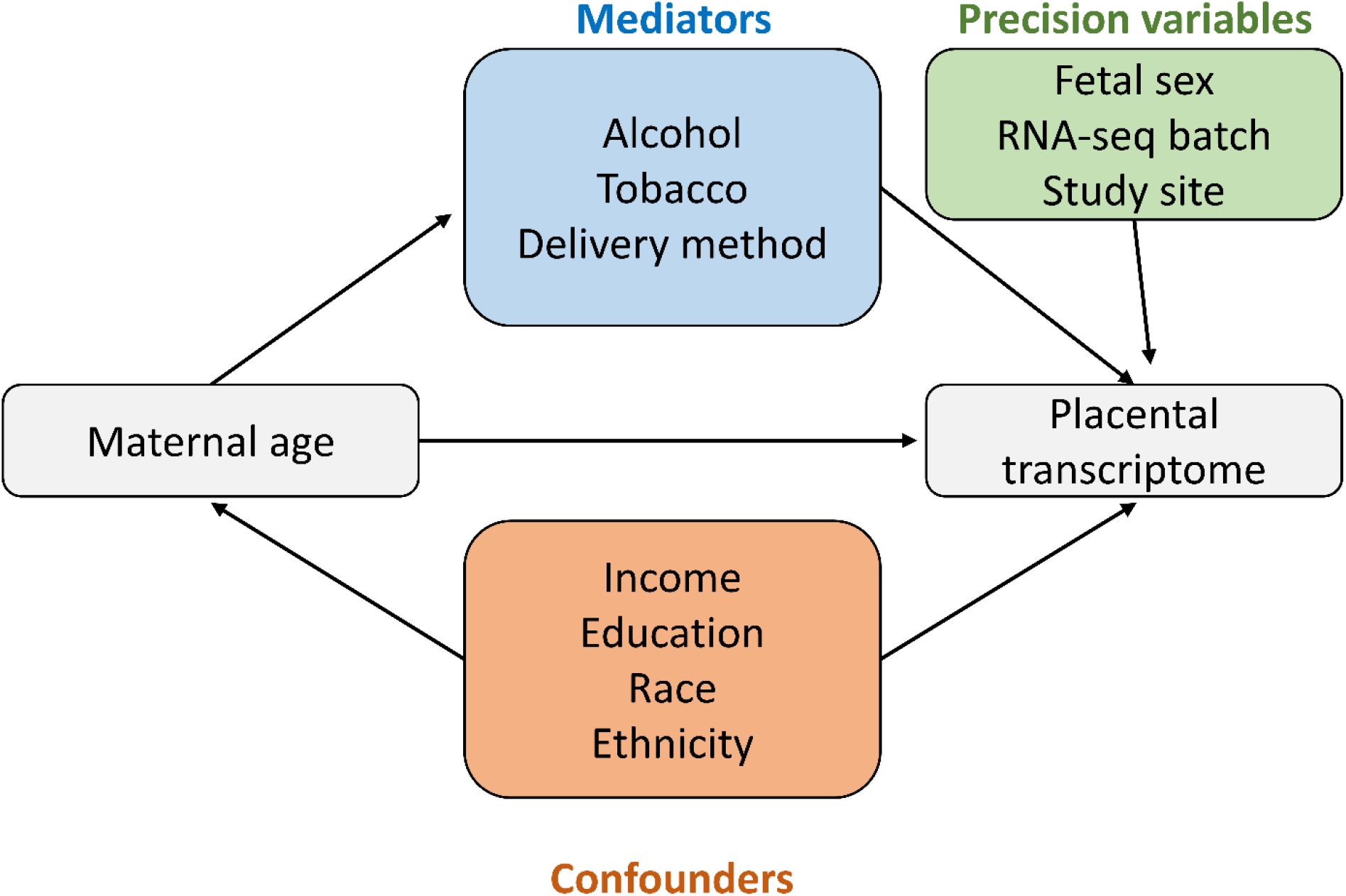
Maternal Age to Placental Transcriptome Directed Acyclic Graph Legend: Conceptual model of association between maternal age (predictor) and the placental transcriptome (outcome). Confounders are upstream causes of both the predictor and outcome. Mediators are on the causal pathway between the predictor and outcome. Precision variables could affect the outcome but have no clear casual effect on the predictor.

Among 1,045 individuals included in this analysis, 6% (61) were missing data for at least one of the 10 covariates. In the CC, SI, and RNAseqCovarImpute MI analyses, maternal age was associated with 575, 214, and 382 DEGs, respectively (Figure 9A). The CC and SI analyses uncovered 96% (368) and 56% (214) of the significant DEGs from the MI method, respectively, while there were 30 DEGs exclusive to MI (Figure 9A). Although there were some differences, genes ranked from lowest to highest P-Value followed similar orders between the methods (Figure 9B). Many of the top DEGs, according to their significance and fold-change magnitude (Figure 9C-E), play roles in inflammatory processes and the immune response. *S100A12* and *S100A8* are pro-inflammatory calcium-, zinc- and copper-binding proteins, *CXCL8* (IL-8) and *IL1R2* are pro-inflammatory cytokines/cytokine receptors, *SAA1* and *CASC19* are known to be expressed in response to inflammation, while *LILRA5*, a leukocyte receptor gene, may play a role in triggering innate immune responses (17).

**Figure 9:**
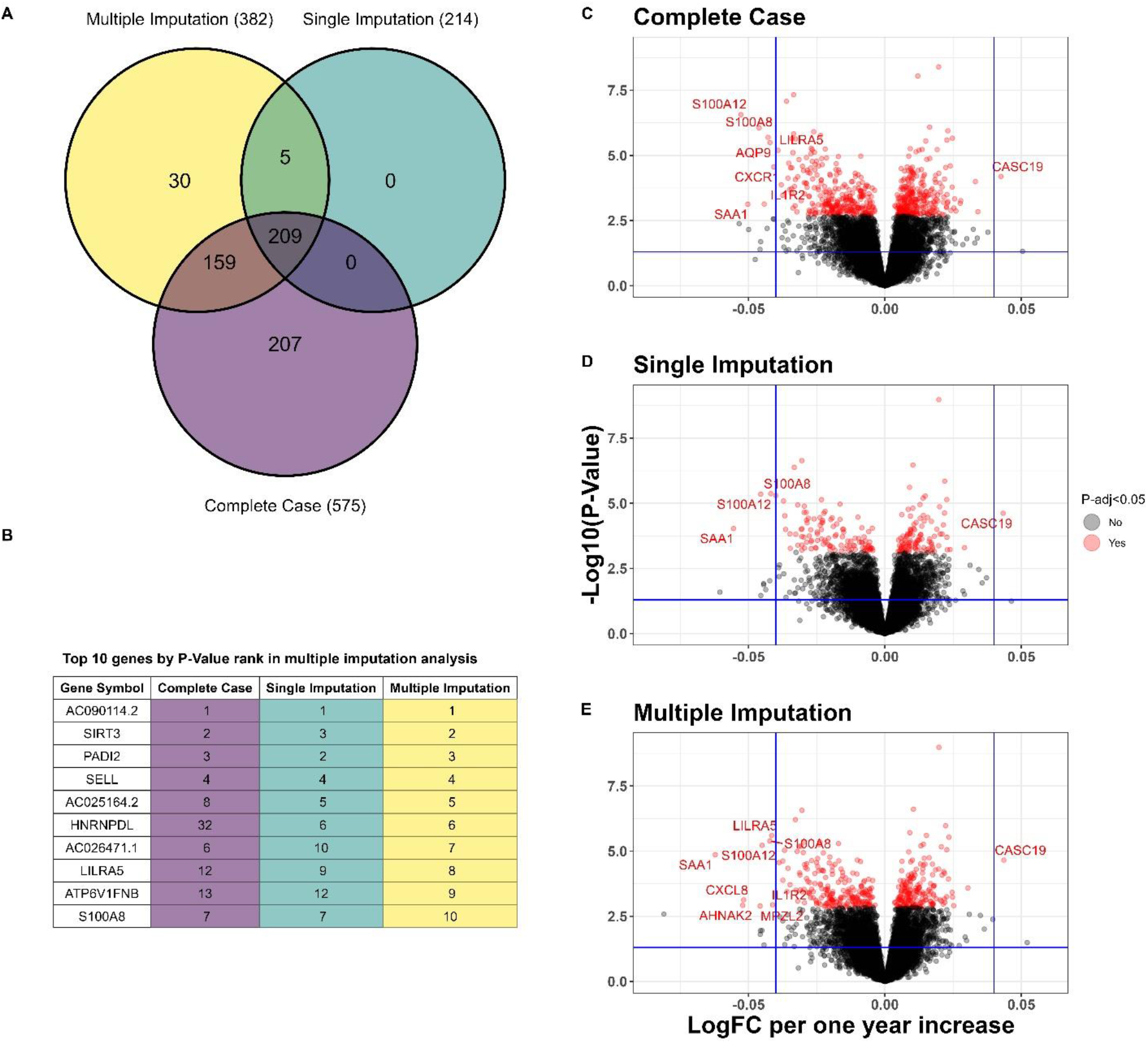
Associations between Maternal Age and the Placental Transcriptome Legend: Venn diagram depicts shared and distinct differentially expressed genes for each method (A). P-Value rankings for each method for the top 10 genes with the lowest P-Values from the multiple imputation analysis (B). Volcano plots of maternal age associations with placental gene expression in complete case (C), single imputation (D) and multiple imputation (E) analyses. Models include the following covariates: maternal race, ethnicity, education, tobacco and alcohol use during pregnancy, household income adjusted for region and inflation, delivery method, fetal sex, sequencing batch, and study site. Log_2_-adjusted fold-changes (LogFCs) shown for each one year increase in maternal age.Horizontal and vertical lines at P = 0.05 and LogFC ± 0.04, respectively. HGNC gene symbols shown for significant genes with false discovery rate adjusted P-value (P-adj) <0.05 and LogFC beyond 0.04 cutoff.

## Discussion

We have shown that a multiple imputation procedure that includes data on the outcome in the imputation predictor matrix has a performance advantage relative to CC and SI methods in RNA-seq studies using the limma-voom pipeline. We found that our newly implemented multiple imputation method in RNAseqCovarImpute was the best performer with respect to identifying true positives, controlling FDR, and limiting bias across a wide range of missing data scenarios. Similar methods allowing for the inclusion of high dimensional outcome data in multiple imputation models have been developed for epigenome wide association studies (EWAS) (8, 9). In EWAS, there appear to be tradeoffs between complete case analysis and multiple imputation, with multiple imputation identifying more true positives but at a higher FDRs (8). In the simulations presented here, however, RNAseqCovarImpute identified more true DEGs with no FDR tradeoff. The application of multiple imputation to EWAS studies represents a more challenging problem owing to higher dimensionality of the methylome (850,000 CpG sites with Illumina’s EPIC array (18)) versus the transcriptome (typically tens of thousands of genes). Moreover, there are additional challenges in applying multiple imputation to alternative epigenomic and transcriptomic methods such as differentially methylated region and pathway enrichment (discussed below) analyses.

To address the sparsity of single-cell RNA-seq data, imputation methods to fill in missing or zero RNA-seq counts have been extensively developed (19). Little attention has been paid, however, to the imputation of missing covariate data in studies where gene expression is the outcome of interest. Lack of development in this area may reflect the fact that a large proportion of RNA-seq studies are conducted in in-vitro or in vivo models and do not suffer from missing covariate data. Complete datasets are common in experimental studies with controlled conditions and a limited number of covariates. In an experimental setting, studies may employ two-group analyses with no additional variables or utilize covariates for which collecting data is trivial (e.g., sequencing batch and sex).

Missing covariate data is ubiquitous, however, in large human observational studies. Methods for the treatment of missing data are well-established in observational epidemiology (20), with multiple imputation increasingly becoming the method-of-choice (21, 22). Yet human observational studies of gene expression have often failed to report on the treatment of missing data, despite its prevalence.

When missing data are explicitly addressed in this context, researchers typically utilize complete case analyses (23-25), while single imputation is a less common alternative (6). Despite its advantages, we are unaware of any studies with transcriptomic outcomes that have utilized multiple imputation for missing covariate data. The simulations presented here suggest that future observational transcriptomic studies may benefit from employing multiple imputation via RNAseqCovarImpute over CC or SI. Moreover, we developed an R package so that users may easily apply the methods presented here to their own data.

We applied RNAseqCovarImpute to a large observation study of maternal age and the placental transcriptome. This analysis assessed the association of maternal age with placental gene expression controlling for confounding variables such as maternal race, ethnicity, and socioeconomic status, and potential mediators such as alcohol and tobacco use during pregnancy. Although there was some overlap, the multiple imputation analysis uncovered a different set of differentially expressed genes compared with the complete case and single imputation analyses. Thus, in a real-world example, the method for dealing with missing data matters, and our simulations suggest that multiple imputation should be the preferred approach.

Nevertheless, any of these missing data methods would be a better alternative than omitting covariate control entirely, a common albeit dissatisfactory approach in observational transcriptomic studies. Failure to control for these variables in analyses of maternal age could lead to erroneous conclusions and even faulty clinical recommendations. For instance, studies have shown that the positive associations of young maternal age with child ADHD in unadjusted analyses are eliminated or even reversed following adjustment for confounding and mediating variables (26-28). Thus, younger pregnancies do not confer increased ADHD risk because of the biology of aging per se but owing to other variables that are correlated with maternal age. Younger mothers are more likely to smoke during pregnancy, and prenatal tobacco exposure may impair neurodevelopment. If the link between younger pregnancy and adverse child development is mediated by increased tobacco exposure, then clinical efforts focusing on reducing tobacco exposures during pregnancy would be more effective than recommendations regarding the ideal age for childbearing (26).

One drawback to RNAseqCovarImpute is that its compatibility with pathway and gene set enrichment methods is currently limited, as many of these methods were developed without multiple imputation in mind. The RNAseqCovarImpute multiple imputation method produces one final list of genes with their associated t-statistics, log fold changes, and P-values for differential expression. Thus, the method is compatible with gene set enrichment analyses that utilize gene rankings such as overrepresentation analysis, or gene level statistics such as camera (29) and gage (30). However, the final gene list produced by RNAseqCovarImpute is based on the combined analyses of multiple gene expression matrices across each imputed dataset. Although theoretically possible, methods that require as input a gene expression matrix or data at the individual sample level are likely not out-of-box compatible with RNAseqCovarImpute. Future work could moderate such methods to accommodate analysis of multiply imputed RNA-seq data.

The RNAseqCovarImpute package is also not out-of-box compatible with all differential expression analysis methods, as it was designed to utilize the limma-voom pipeline. Nevertheless, users may wish to use our multiple imputation procedure with other differential expression methods. The RNAseqCovarImpute package contains separate functions for 1) binning genes, 2) imputing missing covariates, 3) running limma-voom differential expression analysis, and 4) combining results with Rubins’ rules. Thus, although we did not deliberately integrate other methods into our pipeline, the distinct functions for each step allow for the export of the multiply imputed data if users wish to apply alternative expression analysis methods.

## Conclusions

As the cost of sequencing decreases, studies of the transcriptome may experience a substantial shift from small-scale in vitro and in vivo experimental systems to larger-scale clinical and epidemiologic contexts where missing covariate data is prevalent. Multiple imputation is a well-established method to handle missing covariate data in epidemiology, but was previously not compatible with transcriptomic outcome data. We developed and R package, RNAseqCovarImpute, to integrate limma-voom RNA-seq analysis with multiple imputation for missing covariate data, and demonstrated that this method has superior performance compared with single imputation and complete case analyses. Ultimately, RNAseqCovarImpute represents a promising step towards harmonizing transcriptomic and epidemiologic approaches by addressing the critical need to accommodate missing covariate data in RNA-seq studies.

## Materials and Methods

### Multiple imputation and differential expression analysis in RNAseqCovarImpute package Binning genes

The default is approximately 1 gene per 10 individuals in the study, but the user can specify a different ratio. For example, in a study with 500 participants and 10,000 genes, 200 bins of 50 genes would be created using the default ratio. When the total number of genes is not divisible by the bin size, the method flexibly creates bins of two different sizes. For example, if the same hypothetical study included 10,001 genes, 199 bins of 50 and 1 bin of 51 genes would be created. The order of the features (e.g., ENSEMBL gene identifiers) should be randomized before binning.

### Data imputation

Data are imputed using the mice R package with its default predictive modeling methods, which are predictive mean matching, logistic regression, polytomous regression, and proportional odds modeling for continuous, binary, categorical, and unordered variables, respectively (14). The user may specify “M”, the number of imputed datasets, and “maxit”, the number of iterations for each imputation. M imputed datasets are created separately for each gene bin, where the imputation predictor matrix includes all covariates along with the log-CPM for all the genes in a particular bin. Thus, each gene bin contains M sets of imputed data.

### Differential expression analysis

DEGs are determined via the limma-voom pipeline. This procedure fits weighted linear models for each gene that take into account individual-level precision weights based on the mean-variance trend (10). Model results are further moderated with the limma empirical Bayes procedure in which gene-wise variances are squeezed towards a global mean-variance trend curve (11, 12).

Our method first constructs the voom mean-variance curve using all genes within all M imputations. Gene-wise linear models are fit using the lmFit function taking into account the experimental design, with all covariates, separately within each gene bin on each M imputed dataset. The residual standard deviations and average log-counts are extracted from all lmFit models across all M imputations to fit the robust LOWESS curve that is used to estimate voom precision weights (10).

We apply a modified voom function followed by the lmFit function from the limma package (11, 12) separately within each gene bin on each M imputed dataset to estimate precision weights and fit weighted linear models for every gene. We modified the voom function to allow input of bins of outcome genes while utilizing the above precision weights, which were calculated using all genes across all M imputations, rather than estimating precision weights separately within each gene bin. Genes are then un-binned, and the M sets of lmFit model results are stacked into M tables of model results. After un-binning, each M table contains a set of model results for all genes in the analysis. The squeezeVar function from the limma package is then used separately on each M set of lmFit model results to apply the limma empirical Bayes procedure and estimate prior variances and degrees of freedom.

### Pooling results

Rubins’ rules (2) are used to pool coefficients and standard errors, and the Barnard and Rubin adjusted degrees of freedom is calculated (31) (see (3) for more details). Wald statistics are calculated as the pooled coefficients divided by the pooled standard errors, and two-sided P-values are derived from the t-distribution. Rubins’ rules are applied separately for the raw and Bayes moderated lmFit model results. Finally, P-values are FDR-adjusted to account for multiple comparisons (13).

### Real RNA-seq data from ECHO-PATHWAYS

Real placental RNA-seq and covariate data come from the ECHO prenatal and early childhood pathways to health (ECHO-PATHWAYS) consortium (5). This study harmonized extant data from three pregnancy cohorts from diverse populations across the country. The consortium’s core aim is to explore the impact of chemical exposures and psychosocial stressors experienced by the mother during pregnancy on child development, and to assess potential underlying placental mechanisms. To investigate placental mechanisms, the study generated RNA-seq data for the CANDLE and GAPPS pregnancy cohort samples. The generation of placental RNA-seq data for this study is described elsewhere (32). Among the enrolled study sample of 1,503, transcriptomic data are available for 1,083 individuals. We excluded 18 placental abruptions and 20 individuals missing maternal age data, leaving a final sample size of 1,045.

### Simulation study

In the first set of simulations, we used real placental RNA-seq and covariate data from the ECHO-PATHWAYS study. These analyses examined the association of maternal age at birth with placental gene expression while controlling for fetal sex, tobacco use during pregnancy, and RNA-seq batch. We included the study sample described above minus one individual who was missing prenatal tobacco data (N=1,044). The positive tobacco exposure group included individuals with maternal urine cotinine above 200ng/mL (33), as well as individuals who were below this cut-off but self-reported tobacco use during pregnancy. We retained only protein-coding genes, processed pseudogenes, and lncRNAs, and removed genes with average log-CPM<0, resulting in a final sample of 14,027 genes. Log-CPM values were normalized to library size using the weighted trimmed mean of M-values (34). Maternal age was defined as the predictor of interest, while fetal sex, prenatal tobacco exposure, and RNA-seq batch were modeled as covariates.

In the second set of simulations, RNA-seq data were modified to add known signal using the seqgendiff package (16). Rather than create fully synthetic count data from theoretical distributions, we chose this method because it can preserve realistic variability from existing RNA-seq datasets. The method relies on binomial thinning of the RNA-seq count matrix to closely match user defined coefficients. The full data model on the real RNA-seq data contained approximately 82.5% non-significant genes. To match this null association rate, we randomly selected 82.5% of outcome genes and adjusted their maternal age coefficients to approximately 0. Coefficients for the remaining genes were drawn randomly from a gamma distribution. In a sensitivity analyses, we adjusted only 50% of gene coefficients to 0. As described above for the real data, we analyzed the synthetic RNA-seq counts using the limma-voom pipeline and defined significant results as true DEGs before simulating missingness and applying RNAseqCovarImpute, SI, and CC methods. To visualize performance of this method, we plotted our user defined coefficients against the real coefficients produced from the limma-voom analysis on the modified RNA-seq count matrix.

A third set of simulations utilized completely synthetic covariate data with modified RNA-seq counts and explored scenarios with varying starting sample sizes (500, 200, 100, and 50). At each target sample size, we generated five random normal variables (V1-V5) with means of 0 and standard deviations of 1. We defined one variable, V1, as the main predictor of interest. The remaining variables, V2-V5, were defined as confounders of the association between V1 and gene expression. We set correlations between V1 and V2-V5 at Spearman’s rank correlation coefficient of 0.1, but no correlations among V2-V5. For 2,000 randomly sampled genes, we used the seqgendiff package as described above to adjust gene expression associations to match coefficients randomly drawn from a gamma distribution with an 82.5% null association rate. Again, we created full data models using the limma-voom pipeline, simulated missingness, and compared the RNAseqCovarImpute, SI, and CC methods. In a sensitivity analysis, we repeated these simulations at sample size of 200 but with different correlation coefficients (0.2, 0.3, 0.4) between V1 and V2-V5.

### Application of RNAseqCovarImpute in analysis of maternal age and the placental transcriptome

This analysis examined the association of maternal age with the placental transcriptome while controlling for 10 covariates using the ECHO-PATHWAYS sample described above (N=1,045). Covariates included family income adjusted for region and inflation (USD), maternal race (Black vs. Other), maternal ethnicity (Hispanic/Latino vs. not Hispanic/Latino), maternal education (<High School vs. High School completion vs. college or technical school vs. graduate/professional degree), study site, maternal alcohol during pregnancy (yes vs. no), maternal tobacco during pregnancy (yes vs. no), delivery method (vaginal vs. c-section), fetal sex (male vs. female), and RNA-seq batch. The causal relationships among these variables are illustrated in Figure 8. For the maternal race variable, American Indian/Alaska Native, Multiple Race, and Other were collapsed along with White participants to avoid small or zero cell sizes in multivariable models. We retained only protein-coding genes, processed pseudogenes, and lncRNAs, and removed genes with average log-CPM<0, resulting in a final sample of 14,029 genes.

## Code availability

The completely synthetic data and analysis code are available at (https://brennanhilton.github.io/RNAseqCovarImpute_synthetic_data_simulation). The RNAseqCovarImpute package is available on GitHub (for installation instructions see https://brennanhilton.github.io/Impute_Covariate_Data_in_RNA_sequencing_Studies.html).

## Data availability

Real participant data are available upon reasonable request following the data sharing guidelines of the ECHO-PATHWAYS consortium, outlined in LeWinn et al (5).

## Supplementary data

Supplementary materials are available in the file supp.pdf.

