## Supplemental for "RNAseqCovarImpute: a multiple imputation procedure that outperforms complete case and single imputation differential expression analysis"

Supplemental Figure 1:

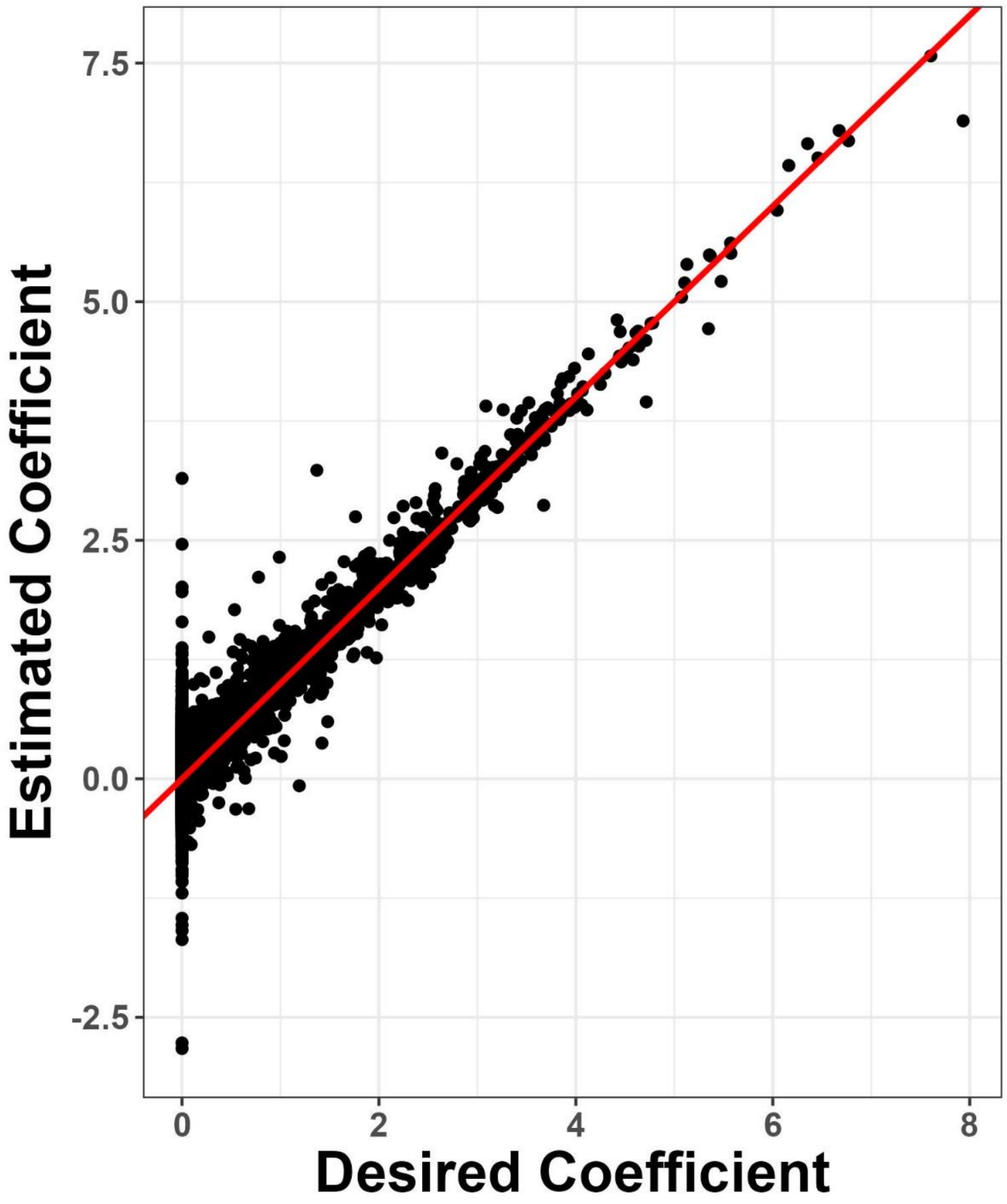


Legend: Desired gene coefficients were input into the seqgendiff package to modify the ECHO-PATHWAYS RNA-seq count matrix. Estimated coefficients come from the limma-voom pipeline applied to this modified count matrix in order to estimate the effect of maternal age on gene expression, controlling for fetal sex, maternal tobacco, and sequencing batch. Data shown from analysis setting 82.5% of gene coefficients to 0.

Supplemental Figure 2:

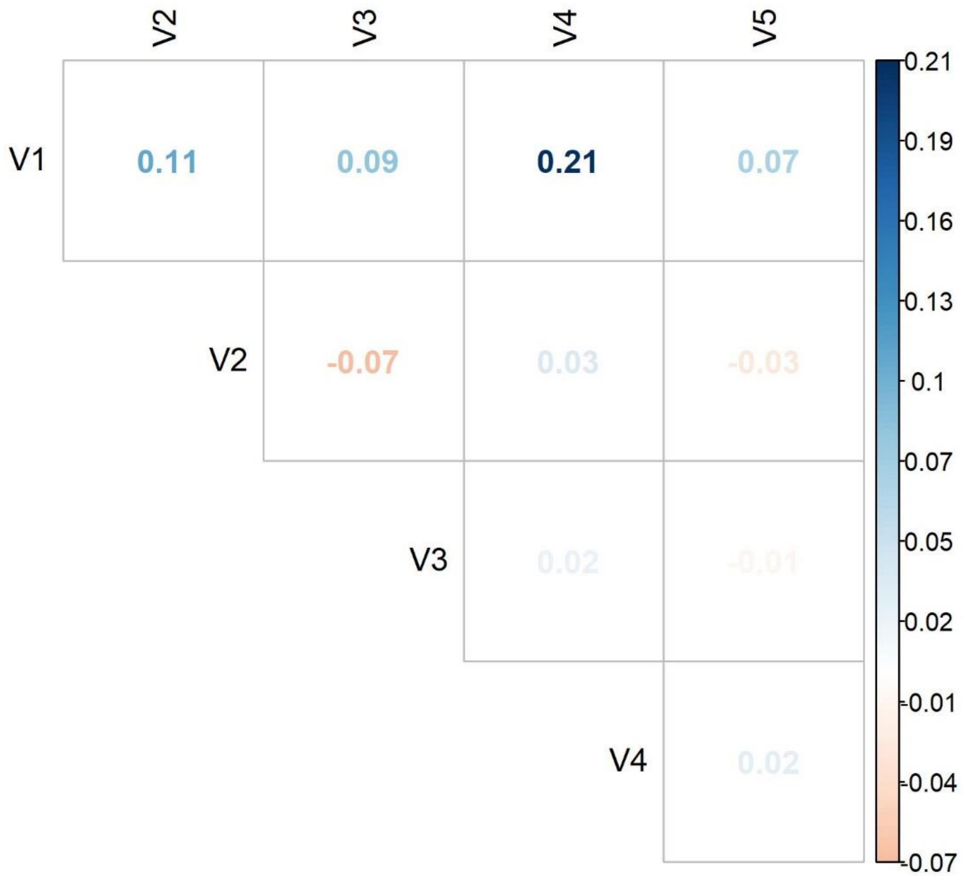


Legend: Correlations among synthetic covariates V1-V5. Synthetic covariates were generated such that V1 (the predictor of interest) was correlated with V2-V5 (confounders) at Spearman's rank correlation coefficient of 0.1.

Supplemental Figure 3:

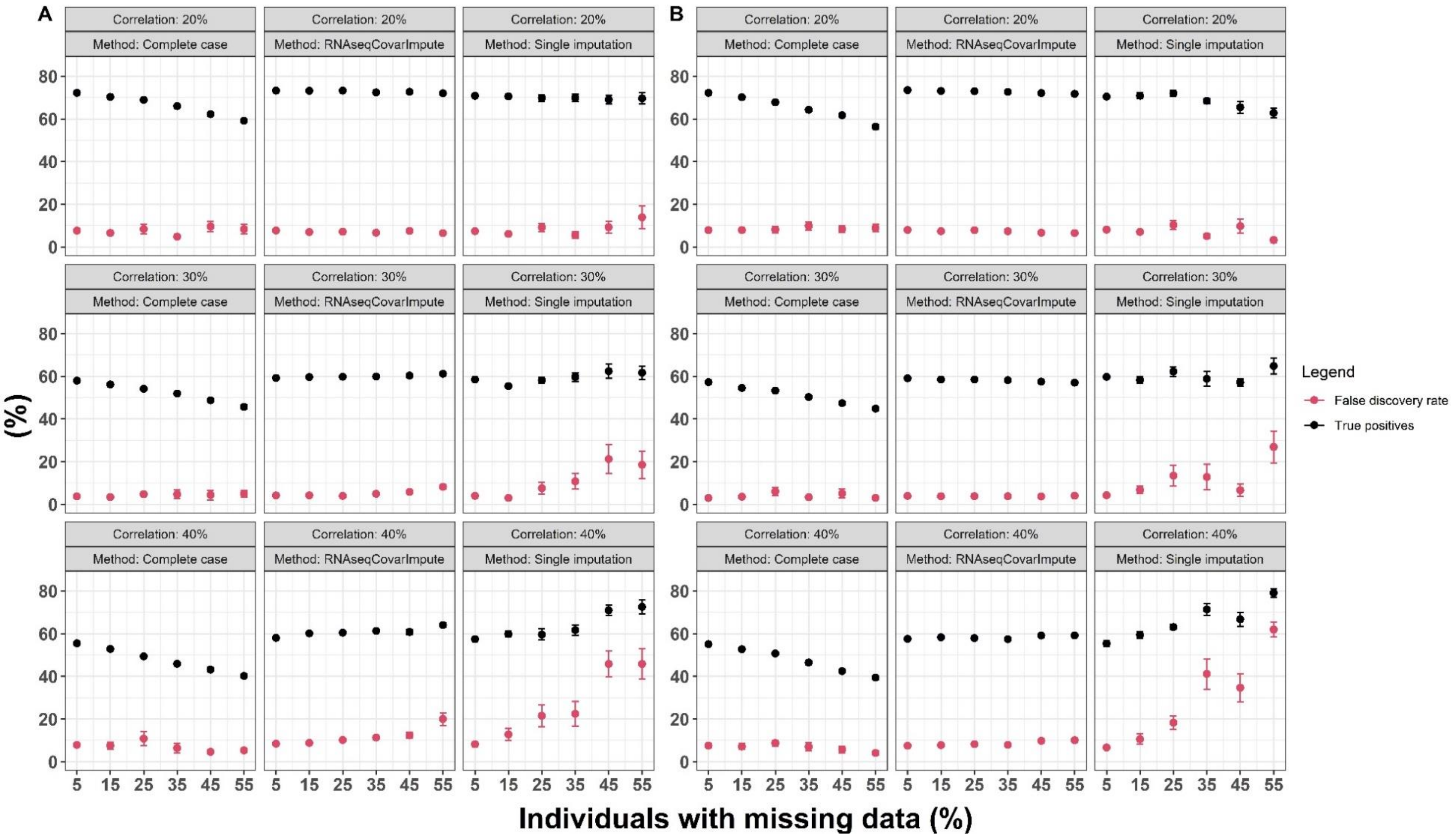


Legend: Performance of complete-case, RNAseqCovarImpute, and single imputation analyses on ten datasets with simulated missingness per missingness mechanism, level of missingness, and correlation level versus the true differential expression model on the fully synthetic covariate and RNA-seq data (N = 200). We set correlations between V1, the main predictor of interest, and V2-V5, confounders, at the Spearman's rank correlations indicated on each panel. Mean ± standard error shown for false discovery rate (red) and percent of true positives identified (black) for MCAR (A) and MAR (B) missingness mechanisms.
